# Comparing phylogenetic trees according to tip label categories

**DOI:** 10.1101/251710

**Authors:** Michelle Kendall, Vegard Eldholm, Caroline Colijn

## Abstract

Trees that illustrate patterns of ancestry and evolution are a central tool in many areas of biology. Comparing evolutionary trees to each other has widespread applications in comparing the evolutionary stories told by different sources of data, assessing the quality of inference methods, and highlighting areas where patterns of ancestry are uncertain. While these tasks are complicated by the fact that trees are high-dimensional structures encoding a large amount of information, there are a number of metrics suitable for comparing evolutionary trees whose tips have the same set of unique labels. There are also metrics for comparing trees where there is no relationship between their labels: in ‘unlabelled’ tree metrics the tree shapes are compared without reference to the tip labels.

In many interesting applications, however, the taxa present in two or more trees are related but not identical, and it is informative to compare the trees whilst retaining information about their tips’ relationships. We present methods for comparing trees whose labels belong to a pre-defined set of categories. The methods include a measure of distance between two such trees, and a measure of concordance between one such tree and a hierarchical classification tree of the unique categories. We demonstrate the intuition of our methods with some toy examples before presenting an analysis of *Mycobacterium tuberculosis* trees, in which we use our methods to quantify the differences between trees built from typing versus sequence data.

---

Phylogenetic trees are a key tool in understanding the relationships and ancestry in a set of taxa. Although evolution often occurs in a way that does not fit a tree model due to horizontal movement of genes, reassortment, recombination, and of course sexual reproduction, trees are used in applications ranging from macroevolution to infectious disease outbreak reconstruction. It is not straightforward to infer phylogenetic trees from data, and sophisticated software packages have been developed for this task [47, 13, 37]. Tree inference typically requires choices between different computational approaches with different underlying prior models, or even different sources of data. In order to understand posterior uncertainty, determine which sources of data are consistent with each other, and ensure that high-quality trees are obtained, it is useful to directly compare reconstructed phylogenetic trees [4, 2, 7, 21, 25].

Existing tools for comparing trees are strictly limited to comparing trees on identical sets of uniquely-labelled tips [50, 41, 42, 4, 48, 33, 25], or trees whose tips are not labelled at all (which may be considered as unlabelled since their labels are unrelated) [36, 34, 29, 8]. There are many applications, however, in which it is important to compare evolutionary trees which have tips with related but not identical labels.

To our knowledge, the mathematical works on developing metrics to address the problem of comparing trees with non-unique labels are [23, 39, 17]. Huber et al. present extensions of labelled tree metrics that require each tree to have the same multi-set for its tip labels, and thus cannot be calculated for trees with varying tip labels. Ren et al. present an extension to the BHV metric [4] which permits distances to be calculated between trees with differing numbers of tips. This measure can be applied to binary trees with a common set of tip labels but where some trees may have ‘additional’ or ‘missing’ taxa. Garba et al. [17] define a probabilistic distance between trees. They note that trees can be compared by pruning them to their set of common taxa, or by an ‘augmentation method’ which uses the union of their tip sets to define the relevant distributions. Where trees have entirely different tip label sets, for example pathogen samples in distinct geographic regions, the former would leave the trees empty and the latter may become unwieldy and fail to draw connections between ‘related’ tips. Our methods enable the comparison of trees whose tips may be partially or entirely different in numbering and labelling, but where there is a natural relationship between tips by membership of ‘categories’, as we now explain.

In Table 1 we summarise some of the many possible applications where trees may have related but not identical tip labels; we describe some of these in more detail in the Discussion section. We refer to the ‘individuals’ which comprise the possible tip labels and the ‘categories’ to which they belong. These categories may also occur as tip labels in other hierarchical classifications or reference trees.

Comparing trees to each other has a wide range of applications in these domains: testing convergence of Bayesian inference methods [28], quantifying differences across different data sets and locations, and using Approximate Bayesian Computation to infer parameters of evolution using simulations, for example of the major bacterial subtypes, or of gene and species trees.

**Table 1.** Examples of applications where individual tree labels may differ but may be comparable according to some broader categories.

| Categories | Individuals |
| --- | --- |
| Bacterial sub-types e.g. serogroups | Bacterial isolates |
| Species | Orthologous genes |
| Hosts | Deep sequencing reads of pathogen from host |
| Protein families | Individual proteins |
| Population groups | Individual organisms |
| Disjoint features or phenotypes | Individual organisms |
| High-level taxonomy | Individual organisms |

Here we describe a distance measure approach to comparing trees whose labels are related at the ‘category-level‘. It is based on capturing trees as vectors which summarise the extent of shared ancestry between taxa. This also leads to a natural measure of ‘concordance’ between trees, which we define. We demonstrate the methods using small example trees to illustrate the intuition. Next we apply the methods to trees inferred from *Mycobacterium tuberculosis* data, demonstrating the power of the method to quantify and identify the differences between trees inferred using MIRU versus whole genome sequence data. Finally, we discuss extensions and further applications of the methods.

## Materials and Methods

### Definitions

In common with much of the literature concerning phylogenetic trees, we will use the following terms:

#### Definition 1.

*A* path *in a tree T is a non-repeating traversal of consecutive edges in the tree. The length of a path is the number of edges on the path. Since there are no cycles in a tree, we know that for each tip i there is a unique path p_i_ from the root to that tip in the tree*.

*The* depth *of a node is the number of edges on the path from the root to that node. The root has depth zero. We require throughout this paper that the root has at least two direct descendants (outdegree 2) in order to avoid spurious apparent dissimilarities between trees which simply have leading edges before the first node with outdegree greater than 1; we prune such edges from the tree*.

*For a pair of tips i and j, their* common ancestors *are the nodes which lie on both paths pi and pj from the root to i and j respectively. The* most recent common ancestor (MRCA) *of a set of tips S is the node with the greatest depth amongst their set of common ancestors. For convenience, we maintain this terminology so that the MRCA of tip i and tip i is the tip i itself. A set of tips S forms a* monophyletic clade *if the MRCA of the tips in S has no descendants that are not in S*.

We propose measures to compare trees where there is a meaningful correspondence between tip labels at the category level but not necessarily at the individual level. Explicitly, let *C* = {*C*_1_,…,*C_n_*} be a set of categories of tip labels, as described in Table 1. For example, each *C_j_* could represent a human patient, and the individual tip labels could be various deep sequencing reads of a pathogen common to all the patients. We will denote the set of all possible individual tip labels as *S*. In this example, *S* could be {`Patient_C1_read1,…, Patient_C1_read_100, Patient_C2_read1,…, Patient_Cn_read_100`}.

Let *σ*: *S* → *C* be the function which associates a tip label with its category, so that if *i* ∈ *S* belongs to category *C_j_* then *σ*(*i*) = *C_j_*. In practice, *σ* can be thought of as a table or data frame, such as:

**Table 2.** Example of a set of categories *C* and all possible individuals *S*, as linked by the function *σ*.

| Category | Individuals |
| --- | --- |
| Patient A | Patient A read 1 |
|  | Patient A read 2 |
| | ⋮ |
|  | Patient A read 100 |
| Patient B | Patient B read 1 |
| | ⋮ |
|  | Patient B read 100 |
| ⋮ | ⋮ |
| Patient N | Patient N read 1 |
| | ⋮ |
|  | Patient N read 100 |

#### Definition 2.

*Let C be a set of categories, S a set of individuals, and σ*: *S* → *C a well-defined (many-one) function which associates each individual with a category. We will refer to a* comparable *set of trees T as a set of rooted trees with associated sets C,S and function σ, where each tree T* ∈ *T has at least one individual tip label associated with each category in C. That is, for each category *C_j_*, T has at least one tip i such that σ(i) = C_j_*.

We note that alternative definitions of tree distances and concordance are possible if this condition is relaxed, but the natural variations we explored placed undesirable weight on some small changes between trees.

We now define a *tip-collapsing* operation * as follows.

#### Definition 3.

*For T* ∈ *T, we define the* collapsed *tree T* to be the tree obtained from T by:*

*1. replacing each tip label i with the label* σ(*i*) = *C_j_*

2. wherever there is a monophyletic clade of tips labelled *C_j_* (tips from the same category) we prune away the whole clade leaving only its MRCA node. This node (which is now a tip) is given the label of the pruned category *C_j_*.

We demonstrate the tip-collapsing operation * with an example in Figure 1.

**Figure 1.**
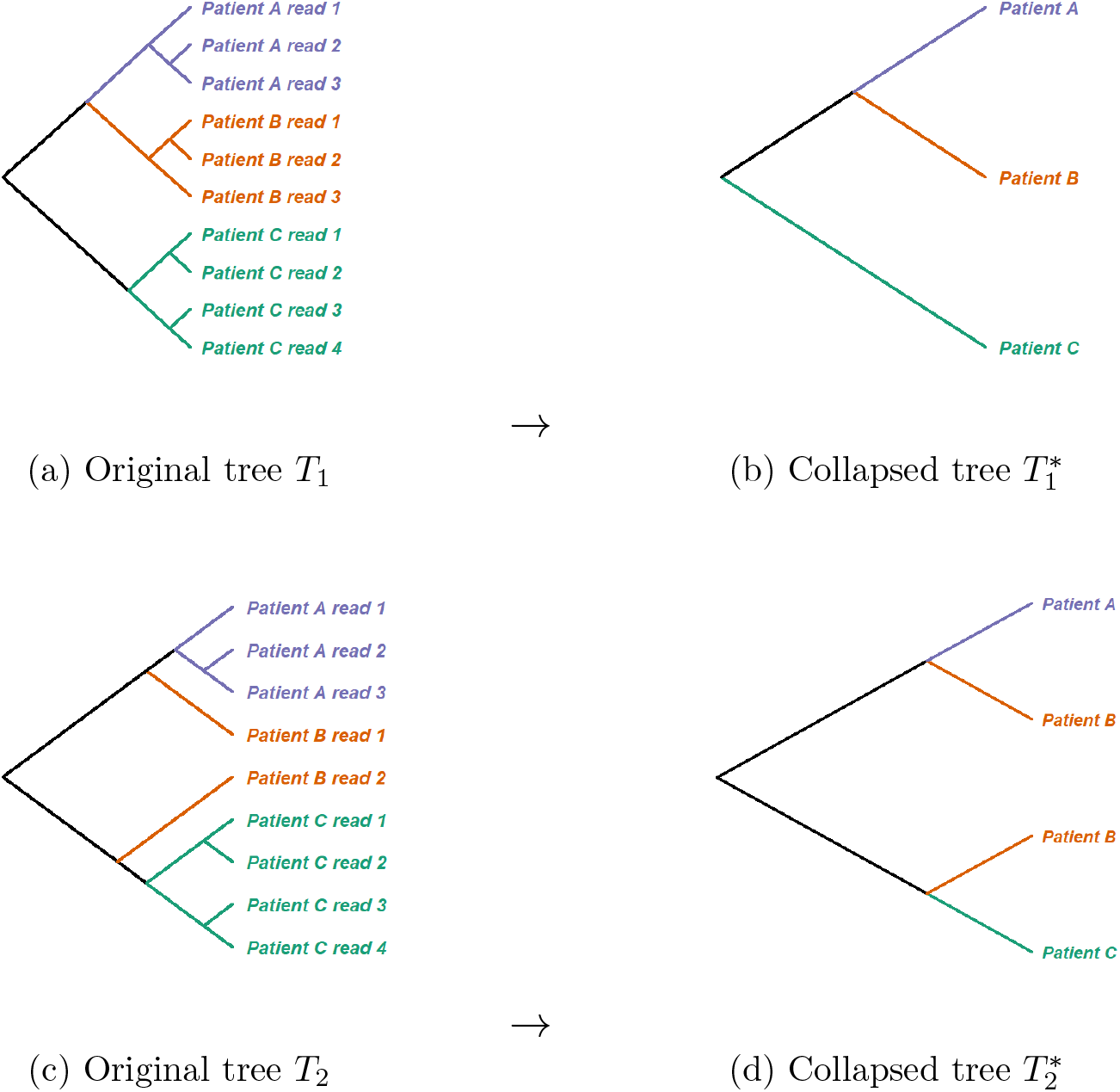
An illustration of the tip collapsing operation. In (b) and (d), each tip from (a) and (c) respectively has been relabelled with its category name (here, the patient ID, also indicated by colour), and each monophyletic clade of same-category tips has been reduced to a single tip with that category label.

### Pairwise tree distance measure

In order to define a distance between a pair of trees we first need to define what it means for two trees to be equivalent.

#### Definition 4.

*We will consider two trees *T*_1_,*T*_2_ in a comparable set of trees P to be* *-isomorphic *if T*_1_^∗^ *and T*_2_^∗^ *have the same labelled shape (topology)* [19, 42], *which is equivalent to saying that there is a tip-label-preserving isomorphism from T*_1_^∗^ *to T*_2_^∗^.

Because the labels of *T*_1_^∗^ and *T*_2_^∗^ may be multisets (some labels repeated as with ‘species B’ in Figure 1), the label-preserving isomorphism may permute multiply-occurring labels. We define a distance measure on the trees *T*^∗^, and by extension, on *T*. In a method similar to those proposed in [6, 25], we characterise each collapsed tree *T^∗^* by a vector *υ*, and then the distance between two trees will be given by the Euclidean distance between their collapsed tree vectors. This distance function is in general not a metric (see below).

#### Definition 5.

Suppose we have a comparable set of trees *P*, with tip labels from *S* and associated *C* and σ. For each tree *T* ∈ *T*, let *P*(*T*) be the set of all tip pairs in T from different categories. That is, if *i, j* are tips in T and σ(*i*) ≠ σ(*j*) then (*i,j*) *E P(T)*. Note that if *T* has exactly one tip associated with each category in C then 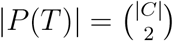.

For a tip pair *(i,j)* ∈ *P(T)*, let *w_ij_(T)* be the depth of the MRCA of *i* and *j* in *T*. (Note that in the case where *T*^∗^ has exactly one label σ(i) and exactly one label σ*(j) then wij(T)* is equivalent to the depth of the MRCA of the category labels σ(*i*) and σ(*j*) in *T^∗^*.) For categories *C_x_* and *C_y_* with *C_x_* ≠ *C_y_*, let *υ_x,y_(T)* be the mean of all the values *wij(T)* for *(i,j)* ∈ *P(T)* where σ(*i*) = *C_x_* and *σ(j)* = *C_y_*.

We can now define the tree vector of T to be

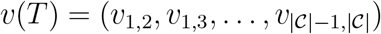

#### Definition 6.

Let ***T*** be a set of comparable trees. For *T*_x_,*T*_y_ ∈ *P* we define the distance between *T*_x_ and *T*_y_ to be:

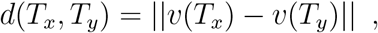

where ║ · ║ is the Euclidean distance (l^2^-norm).

We note some properties of this distance function. Suppose we have a collection of comparable trees *T* = *T*_1_,…,*T_n_, n* ≥ 2, with individual labels *S* and associated *C* and *σ: S → C*. When each collapsed tree *T*_1_^∗^,…,*T_n_^∗^* has exactly one tip label from each category *C_1_*,…,*C_m_* ∈ *C* then *d* is equivalent to the topological tree vector *d_0_* from [25], where it is proven to be a metric (positive, symmetric, zero if and only if the trees are isomorphic, and satisfies the triangle inequality). Therefore, under these conditions *d* is a metric on the collapsed trees *T*_1_^∗^,…,*T*_m_^∗^, and a pseudo-metric on the original trees *T*_1_,…,*T_m_*, which may still differ *within* the structure of their monophyletic category clades even when the distance *d* is zero.

Whenever there are duplicate category labels in any of the collapsed trees *T*_1_^*^,…,*T*_n_^*^, it is possible to find ‘false zeroes’, i.e. a distance of zero between non-isomorphic trees. This is essentially because taking the mean of the depths results in a loss of information. For example, the distance between collapsed trees *T*_1_^*^ = *((A, B), C)* and *T*_2_^*^ = *((A, A, B), C)* (written in Newick format) would be zero. Thus *d* is in general a pseudo-metric on the collapsed trees.

### Concordance measure

The tip collapsing operation also enables us to define a measure of *concordance* between a collection of comparable trees *T* and a reference tree *R*. The labels of the reference tree *R* must also be comparable: they must be the associated set of categories *C*, with the additional condition that each category label must occur precisely once. The concordance value ranges from ≈ 0 if the trees strongly disagree in their evolutionary histories to a value of 1 when they are in ‘complete agreement’, as we explain below.

#### Definition 7.

Let ***T*** be a set of comparable trees and let R be a fixed reference tree whose labels are precisely the associated set of categories C. Extending the earlier notation, let w_x_,_y_(R) be the depth of the (unique) MRCA of categories *C_x_* and *C_y_* in R. Then for any tree T ∈ T we define an indicator for a pair of tips (*i,j*) ∈ *P(T)*:

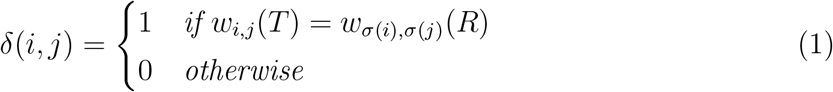

We can now define a measure of concordance between *T* and *R*. It counts the fraction of tip pairs *(i, j) ∈ P(T)* (pairs of individuals in *T* from different categories) whose relative positions in *T* differ from the relative positions of those categories in *R*. Specifically, it counts the fraction of tip pairs for which the depth of the MRCA in tree *T* is the same as the depth of the MRCA of the corresponding categories *σ(i)* and σ*(j)* in *R*.

#### Definition 8.

The concordance of *T* with *R* is given by

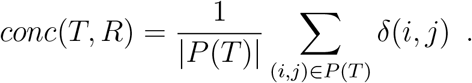

When *conc*(*T*,*R*) = 1, T and R are *fully concordant*, meaning that they describe the same evolutionary relationships at the category level.

#### Theorem 1.

Let ***S*** be a set of individuals and C a set of categories, with an associated function σ: *S* → *C*. Let T be a rooted tree with labels in S and R a rooted tree whose labels are exactly the set C (each occurring exactly once). If conc(T,R)= 1 *then:*

1. if T* is a binary tree, then T* = R. That is, for each category *C_j_*, all its individuals i such that σ(i) = *C_j_* form a monophyletic clade in *T*.

2. if T* is not binary then for each category *C_j_*, all its individuals i such that σ(i) = *C_j_* form one or more monophyletic clades in T, the MRCAs of which are all sisters (they share the same parent node). Thus, if a further collapsing operation were performed on T* to create a tree T**, where any sister tips with the same category labels were collapsed into a single tip with that category label, then T** = R.

*Proof*. Consider the binary case (a). For each category *C_j_*, let the set of individuals *i* such that *σ(i) = C_j_* be denoted *S_j_*. Suppose for a contradiction that *conc(T, R)* = 1 but there exists a set of individuals *S_j_* ⊆ *S* such that *S_j_* does not form a monophyletic clade in *T*. Let *N* be the MRCA of *S_j_* in *T*. Then by definition the direct descendants of *N* are themselves tips from *S_j_* or have descendants which are tips in *S_j_*. Since *S_j_* is not a monophyletic clade, *N* has at least one descendant tip that is not in *S_j_:* call this tip *k*, with σ*(k) ≠ C_j_*. Note that tip *k* cannot be a direct descendant of *N*, otherwise the other direct descendant of *N* would be the MRCA of *S_j_*.

Without loss of generality, let *N*_1_ be the direct descendant of *N* that is an ancestor of *k*, and let *N*_2_ be N’s other direct descendant. Node *N*_1_ has descendants in *S_j_;* call these *S_j_1*, and node *N*_2_ is either a tip itself in *S_j_* or has descendants in *S_j_;* call these tip(s) *S_j_2*. We have that *S_j_1* (J *S_j_2 = S_j_*. The MRCA of any tip *a ∈ S_j_1* and *k* is *N*_1_. In contrast, the MRCA of *k* and any tip *b ∈ S_j_2* is *N*. Since *N_1_* and *N* have different depths, we have *w_k,a_(T) = w_k,b_(T)*. Thus, at most one of *w_k,a_(T),w_k,b_(T)* can be equal to *w_σ(k),C_j__(R)*, hence either *δ(k, a)* or *δ(k, b)* is zero, so *conc(T, R) <* 1.

Now consider the non-binary case (b). Again, assume that *conc(T,R)* = 1 but that there is a set of individuals S_j_ from the same category *C_j_* which is neither monophyletic in T, nor forms multiple sister monophyletic clades in T. Again let *N* be the MRCA of S_j_. Then there exists a direct descendant of N, call it *N*_1_, with at least one descendant tip x in S_j_ and at least one descendant tip k from a different category, i.e. with σ(k) = *C_j_*. Because *N* is the MRCA of all tips in S_j_, it has at least one other descendant y in S_j_ which is not a descendant of *N_1_*. Then the MRCA of x and k has depth greater than or equal to the depth of *N_1_*, whereas the MRCA of k and y is N. As in case (a) this contradicts con(T, R) = 1.

Consider collapsing T to T*. The tips of T* are the MRCA(s) of the monophyletic clade(s) of each S_j_, labelled by the corresponding category *C_j_*. Now, performing the further collapse T**, where any sister tips with the same category labels are collapsed into a single tip with that category label, means that T** and *R* have the same set of tip labels and each occurs precisely once. We then have v(T**) = v(R), which by the Kendall Colijn metric [25] means that T** and *R* are isomorphic.

Finally, we note that whilst the concordance measure can equal one and can reach values close to zero, it will never in fact equal zero.

#### Lemma 1.

The concordance measure conc(R,T) gives values in the interval (0,1].

*Proof*. It is straightforward to construct examples of *R* and *T* such that *conc(R,T)* = 1. It follows from Theorem 1 that for any *T*, letting *T\** = *R* gives *conc(T, R)* = 1.

Suppose for a contradiction that we have comparable, rooted trees *R* and *T* with *conc(R, T)* = 0. Since *R* is rooted we can pick a pair of category labels in *R* whose MRCA is the root (depth 0): call these categories *C_x_* and *C_y_*. Since *conc(R,T)* = 0, the depth of the MRCA of each pair of individual tip labels (i,j) in T with σ(i) = *C_x_* and σ(j) = *C_y_* must be strictly greater than zero, i.e. they are always found on the same ‘side’ of the root in T.

Since the root has outdegree at least 2 (Definition 1), there must be at least one tip in T which descends from a different ‘side’ of the root. Call this tip k and say that it belongs to category *C_z_*, where *C_x_* = *C_z_* = *C_y_*. Now the depth of the MRCA of tip k with any individuals from *S_x_* = {i: σ(i) = *C_x_*} or *S_y_* = {i : σ(i) = *C_y_*} is zero, and so *conc(R, T)* = 0 tells us that the depths of the MRCAs of (*C_x_*, *C_z_*) and (C_y_, *C_z_*) in *R* are both strictly greater than zero. However, we picked *C_x_* and *C_y_* as categories whose MRCA is the root, and tip *C_z_* cannot be on both sides of the root, so we have our contradiction.

We note that requiring every category to be present exactly once in *R* limits the applicability of the method. However, our rationale can be explained by the following simple example. Consider a tree T and let *R* = T*. Then by Theorem 1, *conc(R,T)* = 1. Now consider a tree T′ obtained from T by pruning away all the tips belonging to one of its categories c, along with the internal nodes whose tip descendants all belonged to c. This necessarily removes at least one internal node, with the consequence that at least one remaining internal node will have a shallower depth in T′ than in T. Thus *conc(R, T′)* < 1 despite the remaining tips sharing the same relative phylogenetic *relationships* in *R* and T′ at the category level. Where this is not a concern, the condition could be relaxed.

### Implementation

RAxML trees were inferred using [47]. All other analysis and the presentation of our results were performed using *R* R Core Team 2014. To calculate distances between concatenated WGSs we used the “K80” [26] distance of `dist.dna` from *ape* [31]. The “maximum” and “Euclidean” distances between MIRU types were calculated using dist from the *R stats* package *R* Core Team 2014. The NJ trees were estimated using `nj` from *ape* [31]; UPGMA clustering and ML tree estimation were performed using `upgma` and *pml* respectively from *phangorn* [31, 46]. The MDS of tree distances (Figure 7b) was plotted using `plotGrovesD3` from treespace [24]. Other plotting functions used were from *ape* [31] and *ggplot2* [49].

We have implemented our tree distance and concordance functions in the *R* package treespace [24]. Reproducible code for the small examples in this paper is supplied as a vignette with the package.

### Data

For our analysis of *M. tuberculosis* we chose a dataset representing a cluster of 129 clinical isolates belonging to Lineage 3 that had been characterized both by WGS and MIRU-VNTR typing [3]. A total of 1418 variable sites were concatenated into a single multifasta file by Ayabina et al. and used as input in our analyses.

## Results

### Illustrative examples

We present some small examples to help provide intuition and clarification about the measures we have proposed.

#### Tree distance measure

In Figure 2 we present six comparable trees with individuals set

S = a1, …, a9, b1, …, b9, …, h1, …, h9

where individuals a1, … a9 belong to category A and so on. In Figure 3 we give the matrix of distances between each pair of trees. These figures capture some simple principles behind the behaviour of the distance measure, as we note in the following observations.

**Trees** *T*_1_ and *T*_2_ are very similar, in fact d(*T*_1_,*T*_2_) = 0 because *T*_1_^∗^ = *T*_2_^∗^. The phylogenetic relationships between the categories are identical despite *T*_2_ having more tips than *T*_1_.

**Figure 2.**
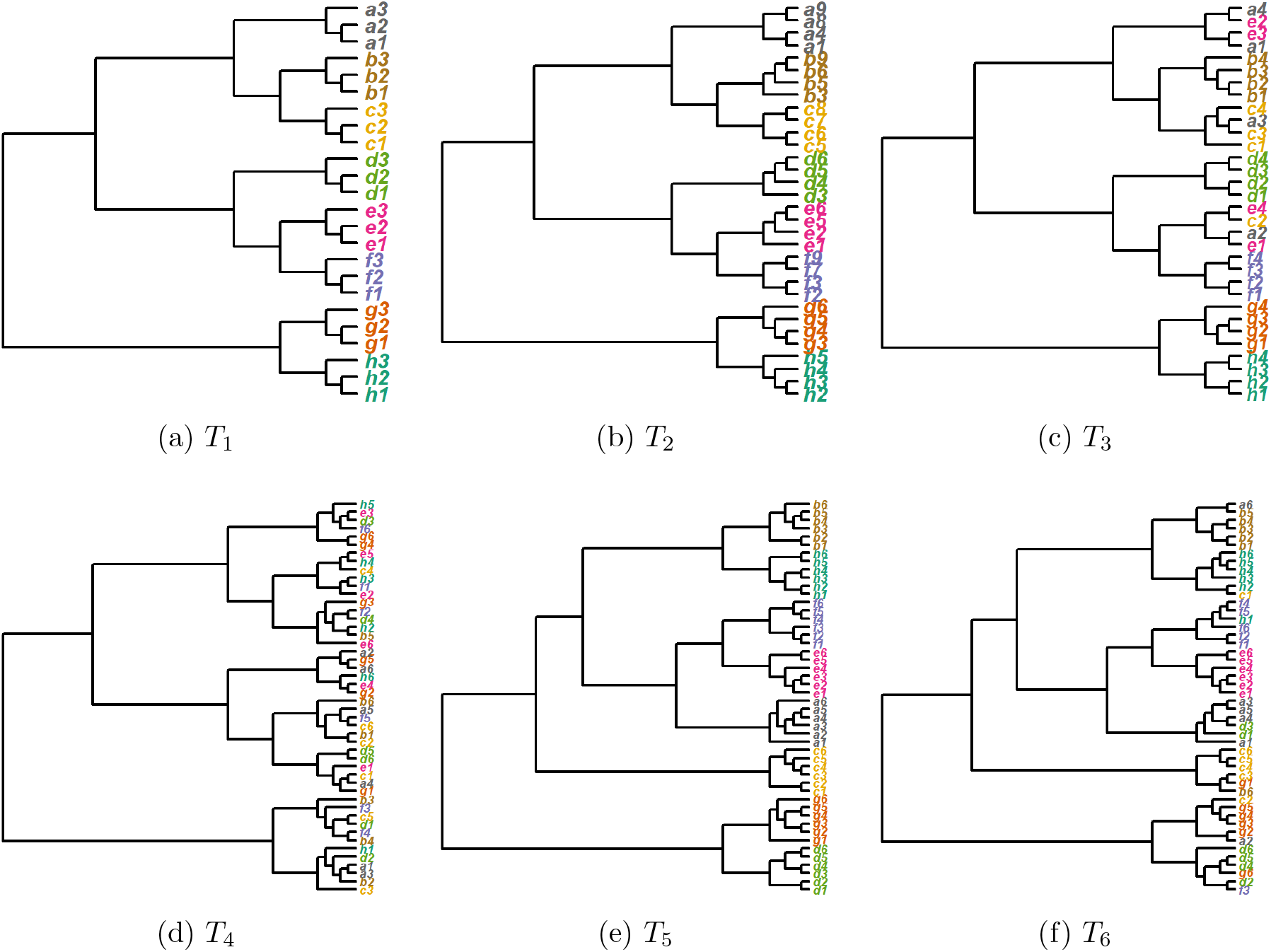
An example of six comparable trees

**Tree** *T*_3_ was generated by taking *T*_1_^∗^ = *T*_2_^∗^, grafting on four individual per category, and then swapping 20% of the tip labels. Accordingly, some of the categories (B,D,F,G,H) remain monophyletic but others are paraphyletic, so that d(*T*_1_, *T*_3_) > 0 (similarly for *T*_2_).

**Tree** *T*_4_ was generated in a similar way to *T*_3_ but by shuffling 100% of the tip labels. None of the categories remain monophyletic, and accordingly d(*T*_1_, *T*_4_) > d(*T*_1_, *T*_3_) (see Figure 3).

**Tree** *T*_5_ was generated by simulating a different, random category tree and appending individual tips to form monophyletic clades. In particular, the pairing of clades for categories *G* and *H* in *T*_1_ and *T*_2_ on one side of the root is replaced by a pairing of category clades *G* and *D* in *T*_5_. We therefore find that there is a rather large distance between *T*_1_ and *T*_5_, since both consist of monophyletic category clades but with different evolutionary histories.

**Tree** *T*_6_ was generated like *T*_5_ but by then shuffling 30% of the tip labels. Accordingly, the distance between trees *T*_5_ and *T*_6_ is quite small (a little larger than d(*T*_1_,*T*_3_), where 20% of tips were shuffled). Notice that the shuffling has created some similarities in category relationships between trees *T*_4_ and *T*_6_, so that d(*T*_4_, *T*_6_) < d(*T*_5_, *T*_6_).

**Figure 3.**
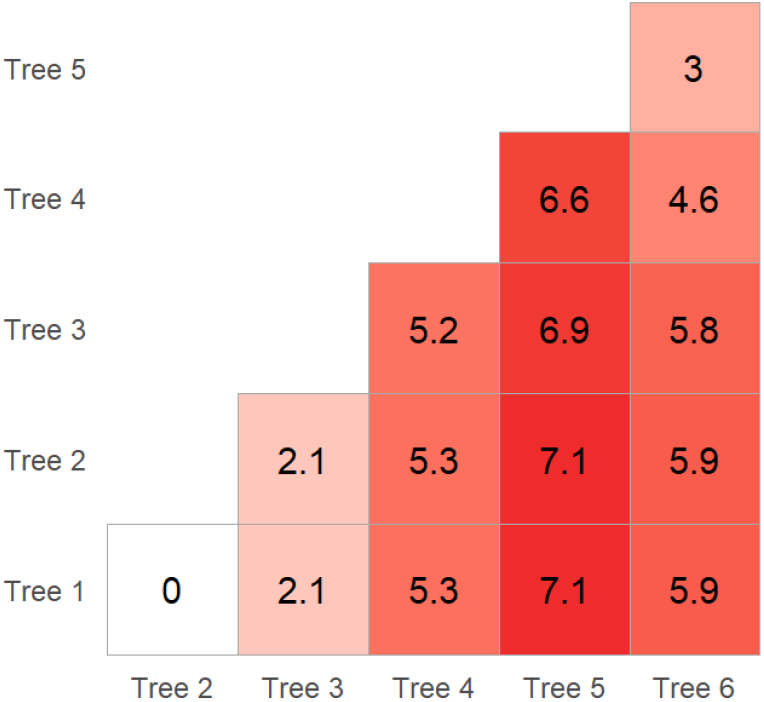
Table of tree distances from Figure 2

#### Concordance measure

In Figure 4 we demonstrate a simple reference tree *R* and three comparable trees *T*_1_,*T*_2_,*T*_3_. We show the concordance of each with *R* in the captions. It can be seen that *T*_1_ is fully concordant with *R* (Figure 4b) because the relationships between the categories are identical: *T*^∗^ = *R*. In *T*_2_ (Figure 4c) the individuals belonging to categories *A* and *B* are no longer each monophyletic, but the overall root separation of categories *A* and *B* from a monophyletic category C is preserved, resulting in fairly high concordance. In *T*_3_ (Figure 4d) the root separates category A individuals from those in categories *B* and *C*, and individuals from *B* and *C* are paraphyletic, resulting in much lower concordance with *R*.

Finally, we demonstrate the effect of ‘mixing’ tip labels in trees so that they are increasingly less concordant with a reference tree. To do this we created a random reference tree *R* with ten tips corresponding to ten categories A to J. To create our comparable trees we then grafted on ten ‘individual’ tips per category. We permuted the labels of either 0%, 20%, 40%, 60%, 80% or 100% of the individual tips, then calculated the concordance of the resulting tree with R. We repeated this experiment 100 times for each proportion of labels to be permuted. Figure 5 is a box plot of the tree concordance results, grouped by the proportion of tip labels permuted. We see that, in general, the more tips that are permuted, the less concordant the tree to the original reference tree R. This reflects the fact that category-level relationships become weaker: as the number of permutations increase, the monophyly of each category gets broken. Some tree similarities remain (and hence the concordance does not decrease below approximately 0.15) due to the fact that each tree is simulated based on the same reference tree R.

**Figure 4.**
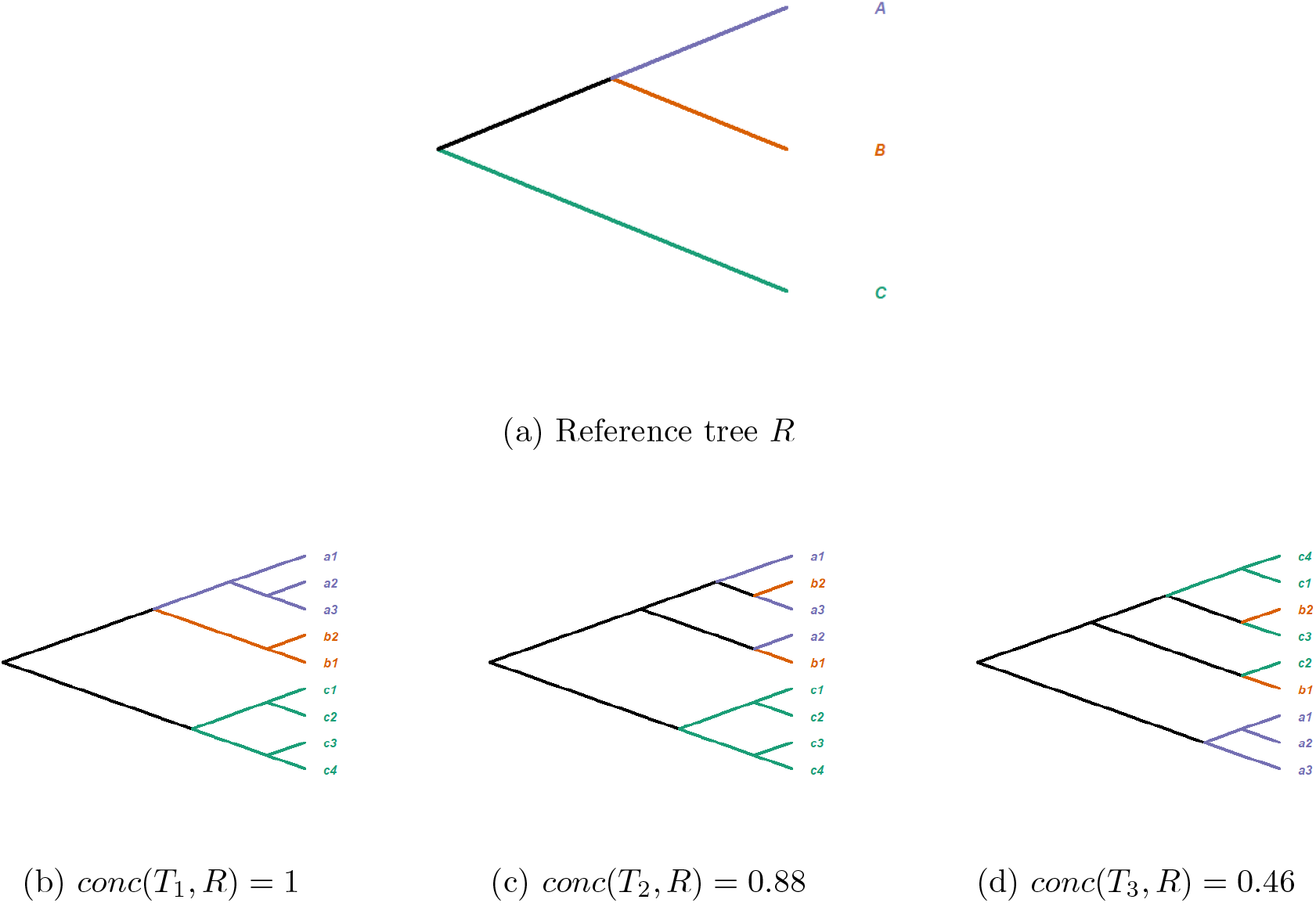
A toy example of a reference tree *R* and three comparable trees *T*_1_, *T*_2_, *T*_3_ with their respective concordance ((c) and (d) rounded to two decimal places).

We note that, as explained by Lemma 1, the concordance does not quite decrease to zero; the minimum value found in this experiment was 0.183111. This tells us that 0.183111 of the individual tip pairs from different categories retained their relative MCRA depths from before the permutation. This number is still relatively high, partly because we did not change the structure of the underlying tree in our experiments. We found that picking a new random ‘base’ tree from which to create our individuals tree each time typically resulted in lower concordance values.

### Comparing MIRU and WGS trees of Mycobacterium tuberculosis

We demonstrate an application of our method to typing and sequence data for the bacterium M. tuberculosis, the causative agent of tuberculosis. Globally, M. tuberculosis remains a leading infectious killer, and molecular epidemiological surveillance is of key importance for identifying and breaking transmission chains. Mycobacterial interspersed repetitive unit-variable number of tandem repeat (MIRU-VNTR) typing is the most commonly applied method for molecular epidemiological surveillance, but whole-genome sequencing (WGS) is rapidly becoming the method of choice in resource-rich countries, due to its superior resolution and power. Since WGS and MIRU-VNTR are very different technologies and MIRU-VNTR samples repetitive regions of the genome that are difficult to capture with short read sequencing, it is natural to expect that trees produced by these input types will differ. This hypothesis is supported by the findings of various studies [5, 43, 14, 18]. Our methods enable us to quantify and visualise such tree differences.

**Figure 5.**
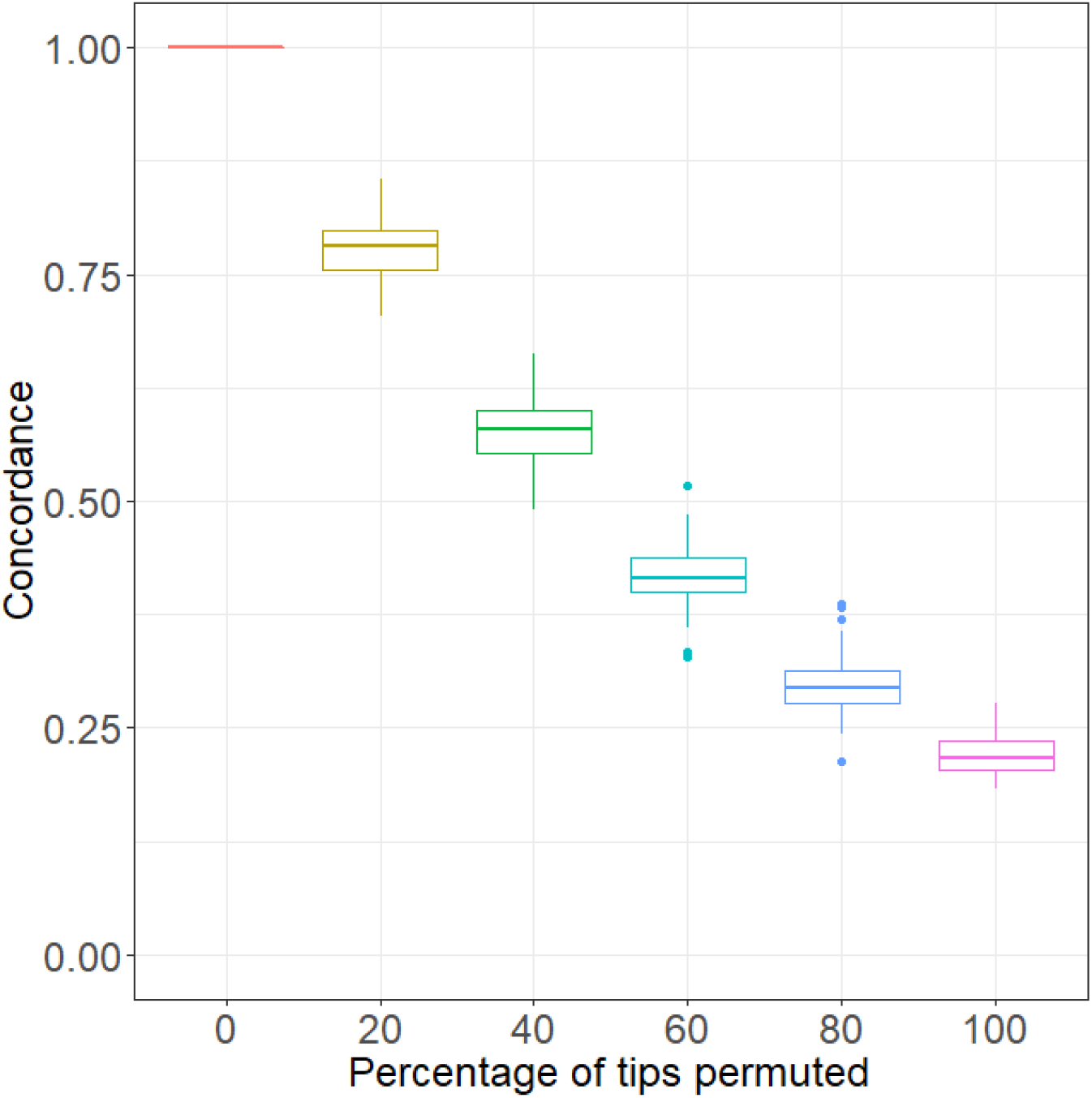
Concordance to a reference tree after a given percentage of random tip permutations

There are a variety of ways to measure distances between whole-genome sequences (see for example the options available for `dist.dna` in the *R* package ape [31]), a variety of measures of distance between MIRU types (including those given in dist in *R* R Core Team 2014), and multiple methods for building phylogenetic trees from a set of distances. We tested as many combinations of these methods as feasible. For simplicity of the narrative we omit here many distances and methods which produced repetitive results. All of our findings and any further combinations can be easily reproduced following the method given in the treespace [24] vignette. Full details of the functions and *R* packages used are given at the end of this section.

We can straightforwardly verify that the differences between isolates according to WGS distances versus MIRU type distances have very low correlation using simple scatterplots, as demonstrated in Figure 6. Here we found the distances between the concatenated whole genome alignment and the “maximum” and “Euclidean” distances between MIRU types.

**Figure 6.**
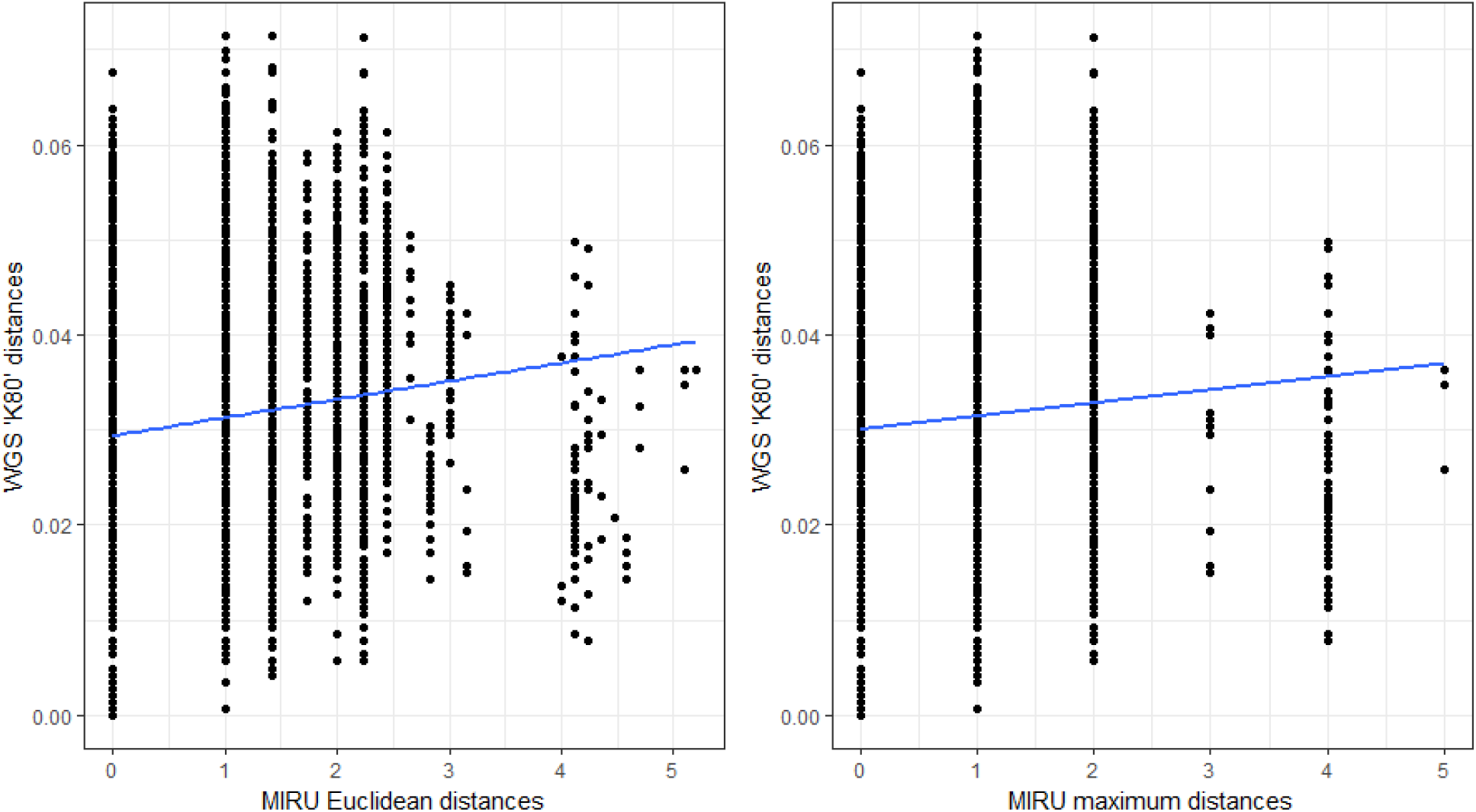
WGS distances and MIRU type distances using the K80 distance for sequences and two choices of MIRU distance.

#### Tree distances

We consider the set of 24 unique MIRU types as ‘categories’, so that each of the 129 isolates can be assigned to a single MIRU category. We can now compare trees at the MIRU category level, even when they have multiple tips per MIRU type.

We made trees of all the isolates (unless stated otherwise) according to the following methods: UPGMA maximum distance on MIRU types; UPGMA Euclidean distance on MIRU types; neighbour-joining (NJ) on WGSs; maximum likelihood (ML) on WGSs in *R* and maximum likelihood using RAxML [47]. For the latter, only the 107 unique WGSs were used, as required by the software.

We rooted the trees using the ancestral MIRU type ‘1064-32’ as an outgroup [3]. For the trees created from MIRU distances this was straightforward because all the tips belonging to this category were monophyletic. However, for the WGS trees this was never the case. Instead, we rooted each of these trees using the largest monophyletic clade of tips belonging to category ‘1064-32‘ as the outgroup. Additionally, we included in our analysis a copy of each of these trees with the remaining tips from category ‘1064-32’ (from smaller clades) pruned out; we see later in the concordance measure that this seems to reduce the bias away from focusing so strongly on the paraphyletic nature of this category. The trees are shown in Figure 7a.

**Figure 7.**
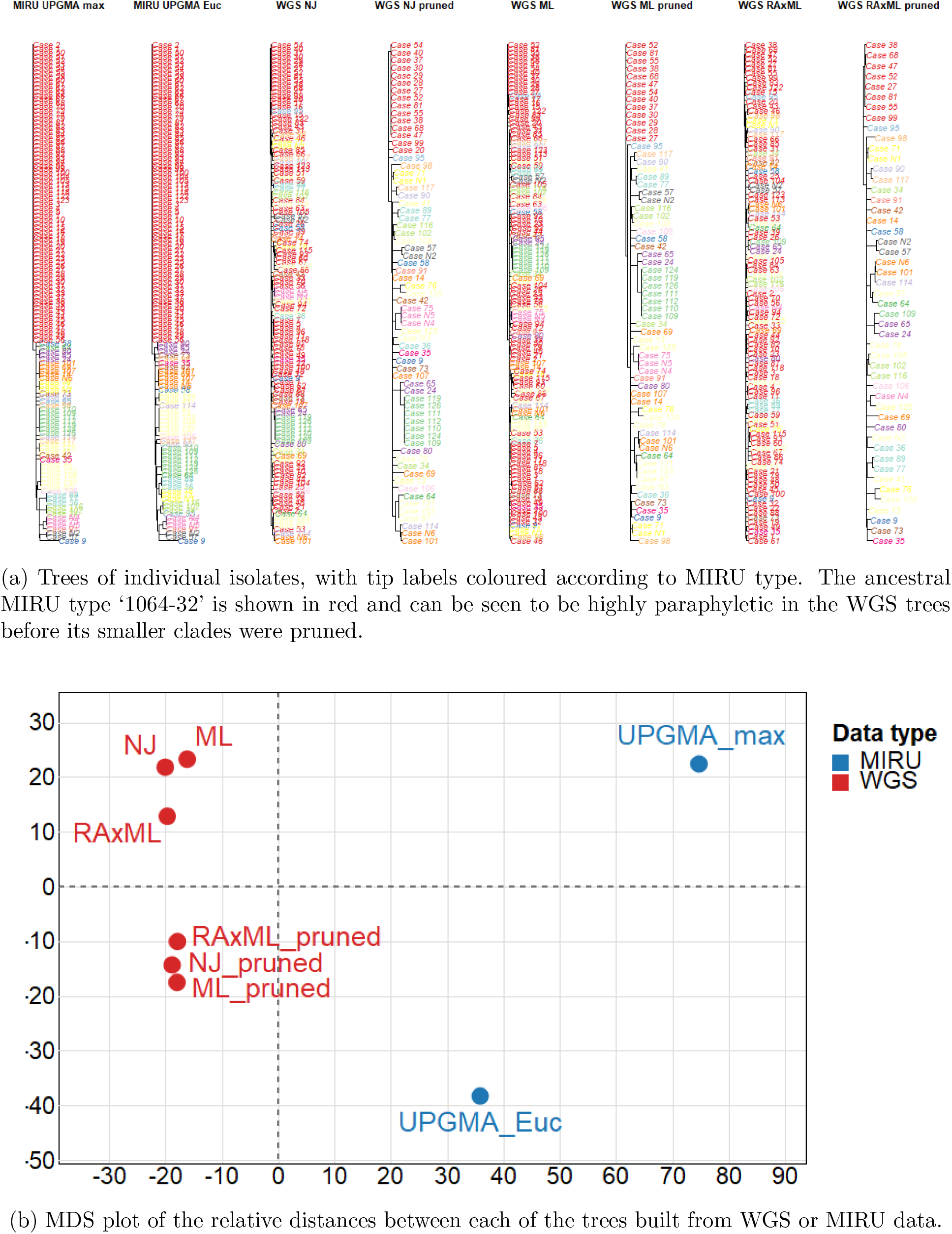
Comparing trees from MIRU and WGS data

Using the distance function (Definition 6) we found the pairwise distances between each of these trees and present them in Figure 7b using a multidimensional scaling (MDS) plot [10]. As in [2, 21, 22, 25], similar trees are plotted closer together, and we can clearly see the difference in signal between MIRU and WGS input data, and the close similarity in the WGS trees after the pruning of non-basal tips of MIRU type ‘1064-32’.

#### Concordance

**Figure 8.**
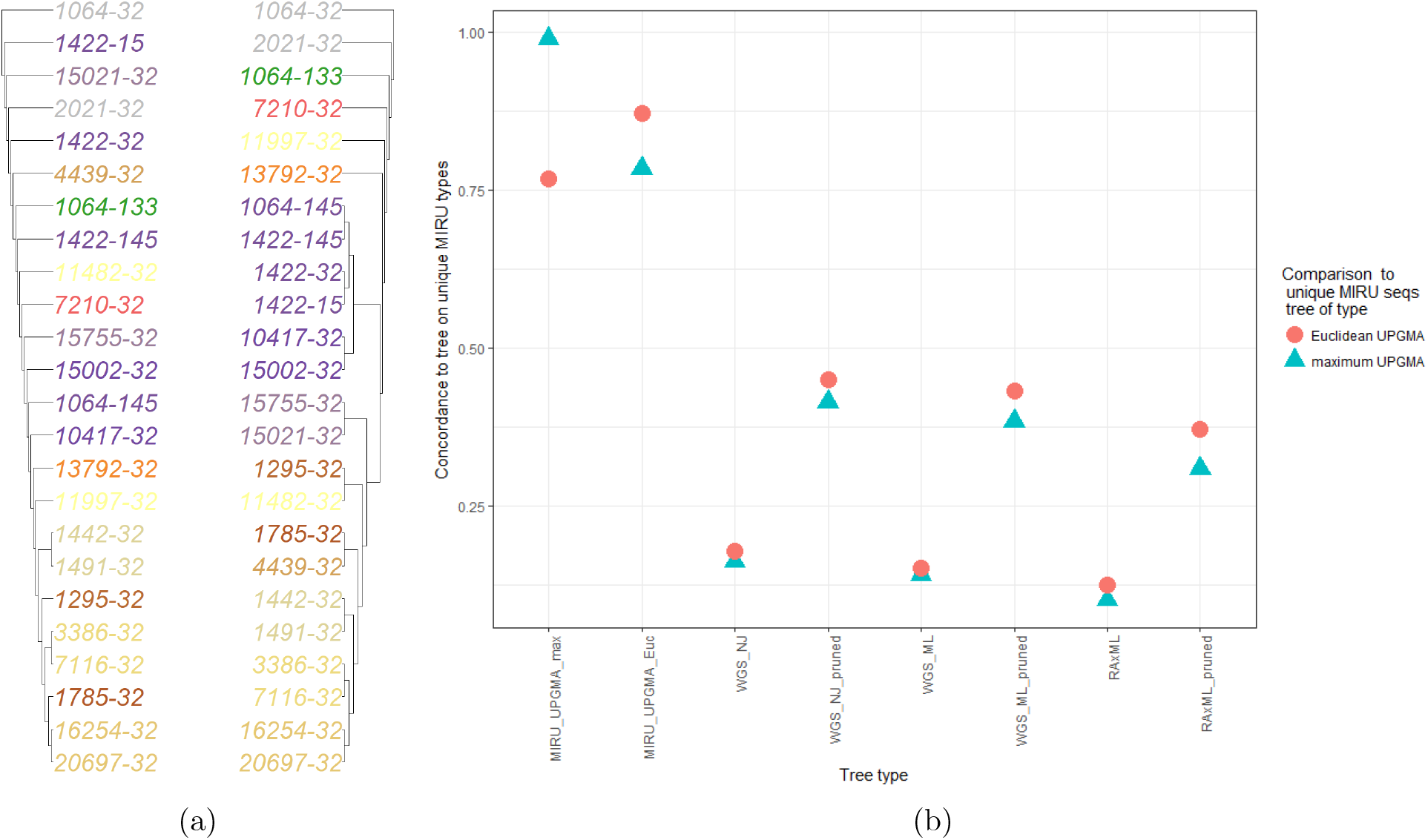
Concordance to MIRU reference trees. The reference trees are given in Figure (a) (“maximum” left and “Euclidean” right), plotted using plotTreeDiff from treespace to highlight topological differences. Tips are coloured according to their number of ancestral differences between the two trees, so tips of the same colour have the same number of differences in their set of MRCA depths between the two trees. Figure (b) shows the concordance of individual isolate trees to the two reference trees.

Using one of each of the 24 unique MIRU types, we built two ‘MIRU reference trees’, one from the the “maximum” and one from the “Euclidean” distance measure and both with UPGMA linkage. These are shown in Figure 8a. We then used the concordance measure to find the concordance of the trees in Figure 7a to these reference trees. The results are plotted in Figure 8b.

The concordance in this setting is a measure of two characteristics of the large trees of individual isolates: first, the extent to which they display MIRU types as monophyletic, and second the extent to which the ancestral relationship between each pair of MIRU types matches the relationship of the corresponding pair in the reference tree. The highest concordances are found between the large trees created from MIRU distances. This was expected because the reference trees were also created from MIRU distances. These MIRU distance large trees did indeed have the property that each category is monophyletic. The highest concordance (0.98) was found between the two trees inferred from ‘maximum’ distance. This reflects the fact that these trees display very similar evolutionary histories per category. The trees from WGS data have noticeably lower concordance, although the concordance is dramatically improved by the pruning of non-basal tips from category ‘1064-32’. We see that, for each large tree, the value of the concordance to the “maximum” reference tree (blue triangles) is quite similar to the value of the concordance to the “Euclidean” reference tree (red circles).

## Discussion

We have presented methods for quantitatively comparing trees according to their heirarchical patterns of a set of categories. We demonstrated one application for *M. tuberculosis*, comparing the phylogenetic signals from MIRU and WGS data. MIRU typing has been noted to be discordant with WGS for tuberculosis previously [5, 43, 14, 18]; our method can assess this quantitatively. This could form a basis for assessing candidate typing schemes, allowing researchers to seek a scheme with the most reliable and high concordance to core genome phylogenies derived from WGS.

Further potential applications are varied and wide-ranging, including new methods for selecting consensus trees [20] and constructing supertrees [45]. In common with the method of Garba et al. [17], the methods also lend themselves to analyses where some or all of the trees have one or more ‘missing’ or ‘additional’ tips due to presence or absence of input data or after the pruning of ‘rogue’ taxa [1, 47]. The methods presented here will place little to no importance on such variations if the ‘missing’ tips (where present) are located close to other tips from their category, and similarly if ‘additional’ tips are grouped close to others from their category. To capture tip presence information, future developments to the methods could include weighting factors to magnify tree distances where there are significant differences between tip sets.

Most bacteria are divided into sub-types according to phenotypic or molecular typing schemes that pre-date next-generation sequencing technologies (examples include the MIRU-VNTR typing we have discussed, as well as MLST typing, capsular typing that defines serogroups of *S. pneumococcus*, sequence types of *S. aureus*, and so on). Consider comparing the evolution of these groups in two geographical areas. In this context it may be important to know how concordant the typing scheme is with a phylogeny (perhaps derived from whole-genome sequencing), how these groups evolved, and whether the loci defining the typing scheme are moving through the population differently than other loci. Datasets from different regions would be expected to mostly share the set of high-level categories, such as serogroups, but will not have the same individual tips. In such applications, therefore, variation in the deep structure of the tree may be of key interest but differences in clades of highly similar taxa from the same category may not be particularly important. Our approach quantifies the concordance at the higher level of categories rather than at the fine-grained level of individual taxa.

At the macroevolutionary level, phylogenetic trees are typically used to represent speciation, with taxa corresponding to species and internal nodes corresponding to inferred speciation events. However, each species contains many genes, and molecular phylogenies derived from gene sequences contain sequences from many orthologous groups of protein-coding genes for each species. Indeed, reconciling gene trees and species trees, and inferring robust species trees from molecular data, is a major challenge for phylogenetics in a macro-evolutionary setting [44, 11, 12, 30]. Nodes in gene family trees may correspond to speciation or to gene duplication, and species’ names occur multiple times in the tips of these trees [9]. Quantitatively comparing gene family trees to each other (for different groups of genes) and to species trees therefore requires coping with trees whose species labels are repeated.

Due to continuing improvements in sequencing technology, it is possible to characterise pathogen diversity within individual hosts. Particularly for HIV, deep sequencing is increasingly used to understand transmission and determine rates of adaptation and immune escape [51, 40]. In deep sequencing studies, different regions of the genome are amplified to produce many reads for each patient, but by the nature of short-read sequencing, reads from one locus are not linked to reads at another. In consequence, while the patient IDs correspond from one locus to another, the individual reads for each patient do not. Applications of deep sequencing are numerous and range from studying the dynamics of immune escape [16, 40], estimating the date of infection [35], comparing diversity in individuals to global diversity, examining reversions to ancestral forms and recombination within individuals [53], and using deep sequencing to infer transmission events [15, 51]. While these tools are most established for HIV, their use in other pathogens, including bacteria, is rapidly becoming established [32, 51, 52].

We note that a single distance between a pair of trees or a concordance value between 0 and 1 can be challenging to interpret. For example, a low concordance value for a tree could be caused by paraphyly of one or more categories, or by significant differences in the relationships between categories in comparison to the reference, or (commonly) a mixture of both. Without plotting the trees it is not straightforward to assess the magnitude of each of these (non-independent) contributions. Another measure or visualisation tool like `plotTreeDiff` from [24] would help to make such results more immediately informative. Similarly, it is not straightforward to define the expected value of either of these measures. Although it may be possible to derive a theoretical expected value (essentially a mean across all possible trees), we would expect trees constrained by data to be much more similar to each other than a pair of random trees. Thus, an objective distance *x* between two trees is not especially informative (unless it is zero) but relative distances *x*_1_, *x*_2_,…,*x*_n_ between n trees can be more readily visualised and interpreted e.g. for identifying clustering or outliers.

The current implementation in *R* is rather slow because it relies on checking for the presence of all possible individual tips supplied in a data frame and repeated vector look-ups. Speeding up the implementation would be desirable and seems likely to be practical.

Finally, we note that our tree distance as we have defined it uses the mean of MRCA depths where there is more than one tip belonging to a given category; other scalar summaries such as the median could also be used. Further, it may be possible to achieve deeper comparisons by retaining the full sets of MRCA positions between tips of one type and those of another, rather than using scalar summaries. Comparing two such sets could be done using the symmetric set difference of the two sets of MRCA depths (in the case that these are discrete) or applying a smoothing operation to continuous depths and comparing sets of depths in the two trees using something like the Kullback-Leibler distance [27]. Additionally, the individual tip-pair contributions could have weighted contributions to the mean (or other summary statistic) according to, for example, the read counts from deep sequencing data, or a measure of importance or credibility on each tip.

